# The Flame Retardant Triphenyl Phosphate Induces Varying Responses in Genetically Diverse Mouse Induced Pluripotent Stem Cells

**DOI:** 10.64898/2026.09.28.754846

**Authors:** Madison Armstrong, Kathryn Janeczko, Hannah B. Dewey, Anne Czechanski, Qiongyu Chen, Emily Swanzey, Callan O’Connor, Whitney Martin, Selcan B. Aydin, Steven C. Munger, Laura G. Reinholdt

## Abstract

Genetic variation impacts biological response to chemical and environmental exposures, contributing to differences in resistance or susceptibility. Forward genetic screens of genetically diverse cell populations can be used to identify the precise genetic variants that drive response variation. Cell lines from laboratory mouse genetic reference populations like the Diversity Outbred (DO) population are well powered for genetic mapping. Diverse stem cell panels are especially beneficial for studying early developmental effects of chemical exposures like triphenyl phosphate (TPHP), an organophosphate flame retardant linked to adverse developmental effects, including altered cell cycle in stem cells. Self-renewal and pluripotency are defining properties of stem cells during development; therefore, it is important to understand how chemicals like TPHP may alter these critical characteristics. To better understand the influence of genetic variation on TPHP exposure response, we utilized a genetically diverse panel of DO induced pluripotent stem cells (DO iPSCs). Our analyses characterize interline variation in gene expression through differential gene expression analysis and expression quantitative trait locus (eQTL) mapping. We identified genomic loci that contribute to variation in gene expression following TPHP exposure, including a regulatory hotspot on chromosome 15. Gene set enrichment highlighted several pathways affected by TPHP including lipid metabolism, steroid hormone signaling, and cell cycle. Our study shows that genetic variation modulates the effects of TPHP on gene expression and cell cycle in stem cells. Additionally, our work demonstrates that genetically diverse cell panels like the DO iPSC resource offer a tractable and scalable approach for identifying the genetic determinants of chemical toxicity.

## INTRODUCTION

An individual’s response to environmental exposure is influenced by interactions between their genetic background and their environment, and exposure response is further affected by developmental stage, tissue or cell type, age, and overall health (Ji and KhuranaHershey 2012; Virolainen et al. 2023). Genetic variation modulates gene expression within these contexts, shaping susceptibility, severity, and ultimately, disease outcomes (Consortiumet al. 2017; Ward et al. 2021). Analogous to mapping causal variants for complex disease phenotypes, gene by environment (GxE) interactions can be used to identify loci that modify adverse-health outcomes associated with environmental exposures (Jayasinghe et al. 2025). Identifying these interactions improves causal inference in toxicology and strengthens chemical risk assessment by directly linking exposures to molecular mechanisms and differential health outcomes.

Flame retardants are a broad class of chemical additives incorporated into consumer and industrial materials to inhibit ignition and reduce the spread of fire. In addition to their fire-suppressive properties, these compounds can add desirable physical characteristics to plastics and films, such as increased flexibility and durability. In recent years, the use of organophosphate flame retardants (OPFRs) has increased following a systemic shift away from legacy toxicants, including brominated flame retardants (BFRs) and plasticizers such as bisphenol A, many of which were restricted or banned in the United States and Europe beginning in 2004 (Ma et al. 2024; Sharkey et al. 2020). Global usage of OPFRs has subsequently expanded, with estimates reaching tens of thousands of tons annually (Chupeauet al. 2020; Ma et al. 2024). OPFRs are now widely incorporated into plastics, furniture, building materials, food-contact materials, personal-care products, and infant products (Chupeau et al.2020; Mendelsohn et al. 2016; Stapleton et al. 2011; Tang et al. 2021). However, many OPFRs are not permanently bound to the materials in which they are incorporated and can disassociate over time, leading to environmental release. Consequently, OPFRs have been detected in surface waters, drinking water, outdoor air, and indoor dust, indicating widespread human and ecological exposure potential (Chupeau et al. 2020; Ma et al. 2024; Marklund et al. 2003;Stapleton et al. 2009).

Among the OPFRs, triphenyl phosphate (TPHP) has come under increasing scrutiny by many of the world’s chemical safety organizations like the United States Environmental Protection Agency (EPA) and Health Canada (Canada 2025; EPA 2016). This attention reflects a growing body of evidence linking TPHP to adverse health outcomes, particularly in reproductive and developmental systems (Mitchell et al. 2018; Shi et al. 2019; Soubry et al.2017). In animal models, TPHP exposure has been linked to neurotoxicity, cardiotoxicity, developmental abnormalities, and tissue disorganization (Baldwin et al. 2017; Qi et al. 2019; Shiet al. 2019). Human epidemiological studies have identified traces of TPHP in human blood, adipose tissue, breast milk, and placental tissues (Chupeau et al. 2020; Hu et al. 2017). A 2017 study investigating the ability of OPFRs to transfer into chorionic villi identified TPHP as one of the most transferrable OPFRs even though it is highly metabolized in maternal tissues (Zhao etal. 2017). TPHP was found to accumulate in fetal tissue during early development due to its slower metabolism in fetal tissue. This accumulation of TPHP and its primary metabolite diphenyl phosphate (DPHP) could result in higher fetal exposure, culminating in potentially more drastic adverse outcomes than those seen in exposed adults (Zhao et al. 2017). Previous studies have identified pathways and proteins affected by TPHP exposure including progesterone synthesis, androgen receptors, peroxisome proliferator-activated receptor gamma (PPARγ), and retinoic acid receptors which all play important roles in development (Bajard et al.2021; Hu et al. 2017). Genes within these pathways harbor genetic variation in human populations and laboratory mice, yet the extent to which this variation modulates individual responses to TPHP exposure is unknown (Chaudhary et al. 2021; Ruth et al. 2016; Skelly et al.2020).

Recent advances in genomics and computational modeling have improved identification of genetic variants that modify response to environmental exposures (Thomas 2010). Translation to human dose-response remains limited by exposure complexity and ethical constraints, particularly for early development. Conventional animal models provide experimental control across life stages but lack genetic diversity, limiting inference on population-level variation (Churchill et al. 2004; Tuttle et al. 2018). Genetic reference populations like the Diversity Outbred (DO) mouse population (Figure 2A) address this limitation by combining experimental control with high allelic diversity. Derived from intercrossing eight inbred founder strains (WSB/EiJ, CAST/EiJ, PWK/PhJ, C57BL/6J, NOD/ShiLtJ, NZO/HlLtJ, A/J, and 129S1/SvlmJ) and maintained by randomized outbreeding, each DO animal carries a mosaic genome that supports high resolution genetic mapping and captures most of the variation present in laboratory mice (Churchill et al. 2012; French et al. 2015; Skelly et al. 2020). However, outbred animal models impose practical constraints. Reproducibility requires large cohorts, and adequately powered studies involve hundreds of animals, which conflicts with animal model reduction goals in the 3Rs framework (Churchill et al. 2012; Gatti et al. 2014;Lauwereyns et al. 2024). To address this, genetically heterogeneous cell lines have been derived from the DO to enable controlled, high-throughput experiments (O’Connor et al. 2024;Swanzey et al. 2021). Building on this approach, our lab generated a panel of DO mouse derived induced pluripotent stem cells (DO iPSCs) to enable controlled, repeatable interrogation of GxE effects (Armstrong et al. 2026). Compared to human iPSC studies, which often have limited allelic diversity, mouse iPSCs provide sufficient genetic variation for genetic mapping and allow validation in genetically matched *in vivo* cohorts (Cortes et al. 2024; Swanzey et al.2021). Using a panel of genetically diverse iPSCs, we found that TPHP exposure alters cell cycle processes and that the magnitude of these effects varies across genetic backgrounds. Transcriptomic analyses identified exposure-response genes and pathways, while genetic mapping revealed loci and genomic hotspots associated with variation in the transcriptional response to TPHP. Together, these results demonstrate that genetic background modulates cellular responses to TPHP and establish the DO iPSC panel as a tractable system for dissecting GxE interactions.

## MATERIALS AND METHODS

### Induced pluripotent stem cell derivation

Tail tip fibroblast lines (P3) derived in a previous study (O’Connor, Keele et al. 2024) were used as starting material for DO inbred founder strains (WSB/EiJ, IMSR_JAX:001145; CAST/EiJ, IMSR_JAX:000928; PWK/PhJ, IMSR_JAX:003715; C57BL/6J, IMSR_JAX:000664; NOD/ShiLtJ, IMSR_JAX:001976; NZO/HlLtJ, IMSR_JAX:002105; A/J, IMSR_JAX:000646; and 129S1/SvlmJ, IMSR_JAX:002448) iPSC reprogramming. Fibroblasts were converted to induced pluripotent stem cells through doxycycline induced, ectopic expression of the four reprogramming genes (‘factors’), *Oct4*, *Sox2*, *Klf4*, and *cMyc* (OKSM) delivered via lentiviral transduction. Lentiviruses were produced from 293T packaging cells following transfection with a 6 plasmid system that included pHage2-EF1a-rtTA-ires-Puro (backbone), phage2-tetO-mOKSM (backbone), pHDM-Hpgm2 (gag/pol), pHDM-tat1b (tat), pRC-CMVRev1b (rev), and pHDM-VsB-G (vsv-g), allowing for the assembly of a single inducible polycistronic cassette as previously described (Mostoslavsky et al. 2006; Sommer et al. 2009). Vectors were a gift from Dr. Matthias Stadtfeld, Weill Cornell Medicine.

Approximately 100,000 DO tail tip fibroblasts were seeded on plastic in 35-mm culture plates (CytoOne) in mouse fibroblast growth media (RPMI 1640 media [Gibco, 11875093] containing 1X penicillin/streptomycin [100X, Gibco 15140], 1X GlutaMAX [100X, Gibco, 35050061], 1X non-essential amino acids [100X, Gibco 11140], 0.0005% 2-mercaptoethanol [55 mM, Gibco 21985023], and 10% fetal bovine serum [Gibco, 16141079]) and infected with 15 μl of concentrated virus in the presence of polybrene (5 μg/ml). Medium was replaced after 16 hours with mouse embryonic stem (ES) cell medium (DMEM supplemented with 15% ES cell grade FBS [Gibco, 16141079], 1X GlutaMAX [100X, Gibco, 35050061], 1X penicillin/streptomycin [100X, Gibco, 15140], 1X nonessential amino acids [100X, Gibco, 11140], 1X sodium pyruvate [100X, Gibco, 11360], 0.1 mM β-mercaptoethanol [55 mM, Gibco 21985023], and 1,000 U/ml leukemia inhibitory factor [Chemicon ESG1106 or 1107], supplemented with 2i [3 μM CHIR99021, Stemgent 04-0004, GSK3 inhibition and 1 μM PD0325901, Stemgent 04-0006, MEK/ERK inhibition]) (Sim et al. 2017) with daily media changes. To initiate reprogramming ascorbic acid was added to stabilize epigenetic marks (Stadtfeld et al. 2012) along with doxycycline (1 ug/ml, Sigma-Aldrich) to induce expression of the reprogramming factors. Doxycycline and ascorbic acid were removed at day 10 postinfection. iPS colonies were collected 20-25 days postinfection and polyclonal cultures were expanded by plating on mitomycin C-treated MEFs in mES cell medium. Cell lines were frozen at P3 in ESM 2i/LIF media containing 10% DMSO and 20% FBS.

Derivation and characterization of the DO iPSC panel is described in our previous resource paper (Armstrong et al. 2026)

### Genotyping

Each of the lines were thawed and expanded to P5-P6 for characterization and banking. Cell culture contaminants were tested using standard bacterial / fungal culture and qPCR for mycoplasma as previously described (Czechanski et al. 2014). DNA was collected from each line and genotyped using the Giga Mouse Universal Genotyping Array (143,000 SNP markers, GigaMUGA; (Morgan et al. 2015)).

We imputed 36-state founder diploid genotype probabilities from SNP-level calls using a hidden Markov model implemented in the *qtl2* R package (Broman et al. 2019). The 36-state diploid genotypes were then condensed to 8-state founder allele probabilities for eQTL mapping (below). Genotype data are available in the Supplemental Files available online.

### Automated High Content Imaging

Frozen aliquots of low passage iPSCs were thawed for each DO founder strain and grown on gelatinized 60mm tissue culture treated plates (CytoOne) in embryonic stem cell media that contained leukemia inhibitory factor (LIF) and inhibitors for GSK-3 and MEK/ERK pathways, also referred to as ESM 2i/LIF. Cells were grown and passaged using 0.05% trypsin-EDTA (Gibco, 25300-054) for at least two full passages to ensure thaw recovery. Collected cells were resuspended in ESM 2i/LIF and counted using a Nexcelom Cellometer Auto T2 Cell Counter. For each strain, working solutions of 12,000 cells/mL were created, followed by plating onto black, gelatinized 96-well plates using the Integra Assist Plus pipetting robot, generating 96-well plates with approximately 1,500 cells per well. Cells were then incubated at 25°C, 5% CO2 for 24 hours. Regular cell media was then swapped for ESM 2i/LIF containing triphenyl phosphate (TPHP; Thermo Fisher Chemicals; dissolved in DMSO) at various concentrations (0, 10, 25, 50, 75, 100, 200, and 400 μM) (Cano-Sancho et al. 2017; Mitchell et al. 2018; Qi et al.2019). Exposure lasted for 24 hours, after which the TPHP media was removed and cells were gently washed with PBS (Gibco, 20012-027) before staining with a cocktail of Hoescht 33342 (Life Technologies, H3570), CellROX Green (Life Technologies, C10444), and Zombie Red dye (BioLegend, 77475) for 20 min. Cells were washed again with PBS and then fixed using 4% paraformaldehyde (PFA; Electron Microscopy Sciences, 15714-S) for 15 min. Fixative was removed and a small amount of PBS was added to each well to prevent the samples from drying out.

Images of inbred DO founder cell lines were captured using the Opera Phenix Plus high content imaging platform equipped with a 20x water/NA 1.0 immersion objective and binning 2. Within each well, multiple planes were imaged, starting at −10.0 μm to 22.0 μm, at 4.0 μm between each plane to quantify whole colony data. Each well provided 57 contiguous images for maximum coverage. Exposure times, focal heights, and excitation power settings for the inbred DO founder screen were Hoechst 33342 (time: 60 ms, power: 40, height: −7.0), Zombie Red (time: 120 ms, power: 80, height: −7.0), CellROX Green (time: 100 ms, power: 100, height: −2.0).

### Image Analysis / Colony Segmentation

Brightfield corrected images were analyzed and processed using Harmony 5.1 software with PhenoLOGIC (PerkinElmer). Stack processing was performed using 3D analysis to quantify whole colony data. Colonies were identified using Hoechst 33342 with absolute thresholds, setting the lowest intensity cutoff at 400, a volume cutoff at > 9000 μm^2^, and with settings for fill and join touching fragments toggled on. Within identified colonies, nuclei segmentation was performed using method “c” within the Harmony software, with a common threshold of 0.06 and a volume cutoff at > 500 μm^2^. Colony morphology properties and number of nuclei were quantified for each colony and well, including mean, median, and standard deviation values were applicable.

### Exposure of iPSCs to TPHP for Dose Response Modeling and RNA-seq

Frozen aliquots of low passage DO iPSC lines were thawed and grown on gelatinized 60mm tissue culture treated plates in ESM 2i/LIF. Cells were grown and passaged using trypsin-EDTA (Gibco, 25300-054) for at least two full passages to ensure thaw recovery. Collected cells were resuspended in ESM 2i/LIF and counted using a Nexcelom Cellometer Auto T2 Cell Counter. Cells were plated onto a 12-well glass-like bottom dish (Cellvis, P12-1.5P) at a cell density of 75,000 cells per well. After 24 hours, the ESM 2i/LIF was replaced with ESM 2i/LIF containing triphenyl phosphate (TPHP; Acros Organics, CAS: 115-86-6; dissolved in DMSO) at various concentrations (0, 10, 25, 50, 75, 100, 200, and 400 μM) for dose-response modeling or 0 μM and 80 μM for the EC20 screen. Cell lines were exposed to TPHP for 24 hours and then trypsinized for cell counting using 0.4% trypan blue (Gibco, 15250-061) as an indicator for cell death. Samples collected for RNA-seq were then pelleted and frozen at −80°C for storage until sequencing.

### Mass Spectrometry

DO iPSC lines were selected based on their percent change in cell count after 24-hours exposure to 80 μM TPHP, representing 2 strong responders, 2 cell lines with close to 20% cell loss, and 2 weak responders, with a male and female line for each response group. Each cell line was split into two wells on a glass-like bottom 12-well dish where one well received 0 μM TPHP containing media (unexposed) while the other received 80 μM TPHP containing media. After 24 hours, media and cell pellets were collected from each well and snap frozen on dry ice for further processing and mass spectrometry analysis. Samples were thawed, treated with 2:2:1 ACN:MeOH:H2O for metabolite extraction. Metabolite supernatants were analyzed using the Thermo Altis Plus QQQ MS via an SRM/MRM assay. Standard curves were generated for TPHP and DPHP in media and cell pellets and used to quantify intracellular and media concentrations of each metabolite per sample (Supplementary Figure 1).

### DMSO Tolerance Testing

Cells from 15 randomly selected DO iPSC lines (7 male, 8 female) were cultured following the same protocol as described above for RNA-seq, except the cells were exposed to 0 μM, 16 μM of DMSO (TPHP vehicle equivalent), or 80 μM TPHP+DMSO. Live cell counts and gene expression data were analyzed and showed no significant effect from DMSO exposure in DO iPSCs at this dose (Supplementary Figure 2).

### Dose-response modeling

The *drc* R package (Ritz et al. 2015) was used to perform dose-response modeling for the rate of cell death in the exposed iPSC lines. For each cell line, we fit four technical replicates to the four-parameter log-logistic dose-response model (Equation 1) using the ‘drm’ function with the ‘fct’ set to ‘LL.4’ and log-normalized cellular features using the ‘bcVal = 0’ option. Model parameters, as shown in Equation 1 include concentration (x), slopes (b), upper asymptotes (d), lower asymptotes (c), and EC50’s (e). Additionally, the ‘ED’ function was used to estimate the EC10, EC20, EC50 for each model fit ‘relative’ to the asymptotes.

Equation 1:

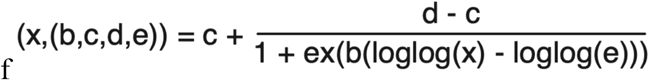

### RNA Sequencing of iPSCs

Total RNA was isolated using the NucleoMag RNA Kit (Macherey-Nagel) and the KingFisher Flex purification system (ThermoFisher). Cells were homogenized in MR1 buffer (Macherey-Nagel) by vortexing, and RNA was isolated according to the manufacturer’s protocol. RNA concentration and quality were assessed using a Nanodrop 8000 spectrophotometer (Thermo Scientific) and the RNA 6000 Pico/RNA Screentape Assay (Agilent Technologies).

Stranded libraries were constructed using the KAPA mRNA HyperPrep Kit (Roche Sequencing and Life Science), according to the manufacturer’s protocol. mRNA was enriched using oligo-dT magnetic beads. Barcoded libraries were created by RNA fragmentation, first and second strand cDNA synthesis, ligation of adapters and barcoding, and PCR amplification. Quality and concentration of the libraries were assessed using the D5000 ScreenTape (Agilent Technologies) and Qubit dsDNA HS Assay (ThermoFisher), respectively, according to the manufacturers’ instructions. Libraries were sequenced 150 bp paired end on an Illumina NovaSeq X Plus using the 10B Reagent Kit and yielded an average of 36 million reads per sample. RNA extraction and QC, library preparation, and sequencing were performed by the Genome Technologies Service at The Jackson Laboratory.

### RNA-seq Processing and Expression Quantification

Raw RNA-seq reads from iPSCs were processed using the Jackson Laboratory Data Science Nextflow DSL2 Workflows (version 24.10.6, https://github.com/TheJacksonLaboratory/jds-nf-workflows/wiki). Reads were aligned and quantified using EMASE (Expectation Maximization for Allele-Specific Expression) against the *Mus musculus* GRCm39 reference genome (Raghupathy et al. 2018). Sequencing quality was assessed using FastQC (Andrews 2010) and summarized with the EMASE pipeline’s integrated QC modules.

Downstream analyses for all RNA-seq datasets were performed in R (version 4.4.0).

Gene expression was normalized per chromosome using upper quartile normalization to account for potential aneuploidy (Evans et al. 2018) and then filtered to select genes expressed in greater than 20% of all samples with a median TPM >= 0.5 in the expressed samples.

Variance component analysis (VCA) was performed to quantify the biological and technical sources that contribute to a proportion of gene expression variability in DO iPSCs. Prior to modeling, tpm-filtered read count values were log-transformed as log10 (read counts + 1) to stabilize variance and reduce the influence of extreme values. Since each genotype is only represented once in each treatment group, we utilized principal components (PC) identified by eigengenes to simulate genotype. Batch refers to the collection date of the iPSCs while run represents the sequencing date. For each gene, we fit the following mixed-effects model:

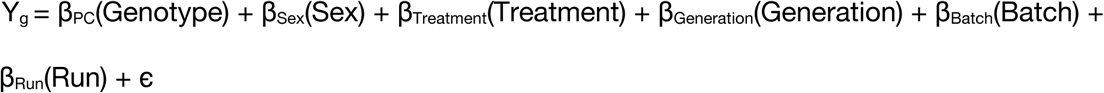

Genes with extremely low overall variance (< 1×10⁻⁵) were excluded prior to modeling, as such genes yield unstable variance estimates. Variance partitioning was carried out using the *variancePartition* R package (Hoffman and Schadt 2016) and visualized using plotVarPart().

### Differential Gene Expression

Differential gene expression analysis was performed using the R package *DESeq2*(Love et al. 2014). We input the expression of 14,430 tpm filtered genes and compared between the unexposed and TPHP exposed populations, using a paired design to account for individual genotype differences. The experimental design accounted for treatment with TPHP, individual genotype, and biological sex implicitly via genotype using the following formula:

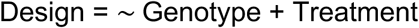

This model estimates main effects of treatment while normalizing expression to each cell line’s baseline. Size factor normalization and dispersion estimation were performed internally by *DESeq2*, followed by fitting negative binomial generalized linear models for each gene.

For exploratory data analysis and visualization, variance-stabilized expression values were generated to place expression values on an approximately log2 scale and reduce the dependence of variance on mean expression. An overall genomic kinship matrix was computed from matched genotype probabilities using calc_kinship (type = “overall”). Eigendecomposition of the kinship matrix was performed and genotype principal component (PC) scores for each sample were calculated. PC scores were standardized prior to downstream use. Variance component analysis (VCA) was performed on the VST-transformed data using the top 500 most variable genes and the top 10 principal components to assess global expression patterns and sample clustering. The percentage of variance explained by each component is reported in Supplementary Table 1.

### Expression Quantitative Trait Mapping and Mediation Analysis

We used the *QTLretrievR* R package (Dewey et al. 2026) to map GxE eQTL. For each expressed gene, we calculated the difference between per chromosome upper quartile normalized counts for exposed and unexposed samples (exposed – control) from each DO iPSC line and then transformed these “delta” values to rank-based inverse normal scores (aka RankZ transformation) to approximate a normal distribution for GxE eQTL mapping. Our mapping model included sex as an additive covariate for the eQTL mapping and identified a LOD threshold for genome-wide significance using 1,000 permutations, establishing a cutoff of LOD > 8.97 for significant delta eQTLs at *α* = 0.05 and LOD > 7.42 for suggestive delta eQTLs at *α* = 0.4.

Delta eQTLs that mapped within ±2Mb of their target gene were classified as local eQTLs, while those mapping outside this window were classified as distant eQTLs. Hotspots of distant eQTLs were identified by summing the number of suggestive or greater distant eQTL (LOD > 7.42) in sliding windows of 50 chromosome-based markers (10 marker shift) across the genome. The top 0.5% of bins having the most eQTLs were considered as hotspots.

We performed mediation analysis on the delta eQTL hotspots to identify potential regulators of the hotspot target genes using the *QTLretrievR* R package (Dewey et al. 2026). We utilized the “double-lod-diff” method to reduce the effects of missing expression values. Mediation analysis was performed on all significant and suggestive and greater distant eQTLs (LOD>7.42) and the top five performing mediators for each distant eQTL were recorded.

### Gene Set Enrichment and Overrepresentation Analysis

Gene set enrichment analyses were performed in R using *fgsea* (Korotkevich et al.2021) on the differentially expressed genes identified by *DESeq2* and all delta eQTL target genes, which were identified using the direct gene connected to each eQTL peak. We defined the universal background to include all genes expressed in our DO iPSCs that passed the tpm and per chromosome mean count normalization described above (14,430 genes).

Manual curation of cell cycle and cell cycle sub mechanism gene sets utilized MSigDB C5 ontology gene sets. Cell cycle was defined by “cell cycle | cell_cycle” while the sub mechanisms were defined as follows:

DNA repair/replication: “dna_repair | dna_replication | double_strand_break | mismatch_repair | replication_fork | genome_stability”

Spindle/chromosome segregation: “mitotic_spindle | spindle_assembly | kinetochore | chromosome_segregation | sister_chromatid | cytokinesis”

G1/S checkpoint: “g1_s_transition | cell_cycle_checkpoint | cyclin_dependent | g1_arrest”

Overrepresentation analyses were performed on CTD-eQTL overlap genes and CTD-TPHP gene set using the *clusterProfiler* package in R (Xu et al. 2024). Delta eQTL target gene human orthologs were identified by the homolgene function in R, using the mouse annotation package *org.Mm.eg.db*. After using the human orthologs to filter for the overlapping CTD-eQTL genes, the overrepresentation analysis was performed using the mouse Ensembl gene ID for consistent use of the mouse MSigDB. For both CTD-eQTL overlap analyses, including the cell cycle enrichment, the universal background was set to include all genes expressed across our control and treated DO iPSCs that passed the tpm and per chromosome mean count normalization described above (14,430 genes). The universal background we defined for the full CTD-TPHP gene list, the CTD-TPHP cell cycle enrichment, and permutation testing for null odds ratio was all genes annotated in the CTD across all species and chemicals. The permutation test included 1000 permutations to identify the threshold for odds ratio cutoff (*OR* 95th percentile threshold = 8.4 across 40 control terms) for gene sets taken at random from the full CTD gene list of similar size to our manually curated cell cycle gene set.

### Flow Cytometry

Frozen aliquots of low passage inbred DO founder iPSCs were thawed and grown on gelatinized 60mm tissue culture treated plates in ESM 2i/LIF. Cells were grown and passaged using trypsin-EDTA (Gibco, 25300-054) for at least two full passages to ensure thaw recovery. Collected cells were resuspended in ESM 2i/LIF and counted using a Nexcelom Cellometer Auto T2 Cell Counter. Cells were plated onto a 12-well glass-like bottom dish (Cellvis, P12-1.5P) at a cell density of 100,000 cells per well. After 24 hours, the ESM 2i/LIF was replaced with ESM 2i/LIF containing triphenyl phosphate (TPHP; Acros Organics; dissolved in DMSO) at concentrations of either 0 μM or 60 μM (inbred DO founder EC20). Cell lines were exposed to TPHP for 24 hours and then trypsinized for collection. Collected cells were washed with PBS and then fixed in ice cold 70 % ethanol at 4°C for 30 min. Cell pellets were isolated and transferred to FACS tubes with 200uL propidium iodide to stain DNA content, samples were then incubated at room temp in the dark for 15 min. Quantification of DNA content was obtained using a BD FACSymphony A5. Cell cycle characterization was performed in FlowJo (10.10.1) using a Watson (pragmatic) model.

### Software

All analyses were performed in R version 4.4.0 (R Foundation for Statistical Computing) on Ubuntu 22.04.4 LTS. Gene expression and related analyses were conducted using Bioconductor packages including *org.Mm.eg.db* (3.20.0), *org.Hs.eg.db* (3.20.0), *biomaRt* (2.62.1), *homologene* (1.4.68.19.3.27), and *variancePartition* (1.36.3). eQTL mapping was performed using the *QTLretrievR* package (1.2.0.0). Additional data processing was conducted using *dplyr* (1.1.4), *tidyr* (1.3.1), *readr* (2.1.5), and *matrixStats* (1.5.0). Visualization was implemented using *ggplot2* (3.5.1), *ggrepel* (0.9.5), and *pheatmap* (1.0.12). RNA-seq processing and quantification were performed using the Jackson Laboratory Data Science Nextflow DSL2 workflows (https://github.com/TheJacksonLaboratory/jds-nf-workflows, version 24.10.6), with alignment and quantification carried out using EMASE. Figures were assembled using Adobe Illustrator.

## RESULTS

Triphenyl phosphate (TPHP) induces cytotoxic effects in mouse pluripotent stem cells, although most prior studies have evaluated these responses in a single genetic background which limits inferences about heritable variation in sensitivity (Qi et al. 2019). While data regarding chemical uptake and clearance rates within stem cells is generally limited, it has been shown that TPHP is taken up quickly by stem cells in culture, peaking within hours of the initial exposure and clearing over the subsequent 48 hours (Qi et al. 2019). Pluripotent stem cells are known to have comparatively low expression of xenobiotic metabolizing enzymes (Kondo et al.2014). To confirm and characterize genetic background differences in cellular uptake, we quantified intra- and extra-cellular TPHP concentrations within a subset of our DO iPSCs using mass spectrometry. Our data demonstrate cellular uptake of TPHP in DO iPSCs with significant difference between genetic backgrounds (Supplementary Figure 1). We were unable to determine if DO iPSCs metabolize TPHP into its diphenyl phosphate (DPHP) metabolite since TPHP spontaneously converts into DPHP in aqueous solutions (i.e. cell culture media) (Supplementary Figure 1). We found that the concentration of DMSO used in our TPHP solution does not significantly affect cell survival or gene expression at our working dose (Supplementary Figure 2).

To determine whether response to TPHP exposure differs across genetic backgrounds, we utilized induced pluripotent stem cell (iPSC) lines derived from the eight inbred founder strains of the Diversity Outbred (DO) population (Figure 1A). These inbred iPSC lines were exposed to a series of eight increasing TPHP concentrations to characterize dose-response relationships and identify heritable variation in exposure-related traits. Previous *in vitro* studies found TPHP to induce oxidative stress and DNA damage, increase apoptosis, and inhibit proliferation and differentiation (Qi et al. 2019; Xiong et al. 2023; Xu et al. 2025). For our study, we used high content imaging to quantify the number of nucleated cells, oxidative stress, cell prominent cellular phenotype that differed significantly among the founder strains was colony volume. Colony volume decreased with increasing TPHP concentration relative to unexposed controls across all founder cell lines, although the magnitude of the concentration-dependent response varied by genetic background (Figure 1C). Notably, the colony volume of cell lines from the more recently wild-derived strains CAST/EiJ, PWK/PhJ, and WSB/EiJ was more resilient to TPHP exposure compared to A/J, C57BL/6J, 129S1/SvImJ, NOD/ShiLtJ, and NZO/HlLtJ (Figure 1C). To further compare these varying responses, we fit a four-parameter log-logistic dose-response model to live cell count data and evaluated the resulting dose-response parameters (Figure 1D). While all cell lines were plated at the same initial density, cell counts in untreated samples differed across genetic backgrounds after 24 hours of culture (Figure 1D). This recapitulates previously published data regarding genetic influence on proliferation rates in mouse embryonic stem cell (mESC) lines from these strains (Skelly et al.2020). In addition to the baseline differences in colony volume, the rate of change in cell count following TPHP exposure differed between the inbred founder iPSC lines (Figure 1E), suggesting that genetic background influences TPHP exposure response in pluripotent stem cells.

**Figure 1.**
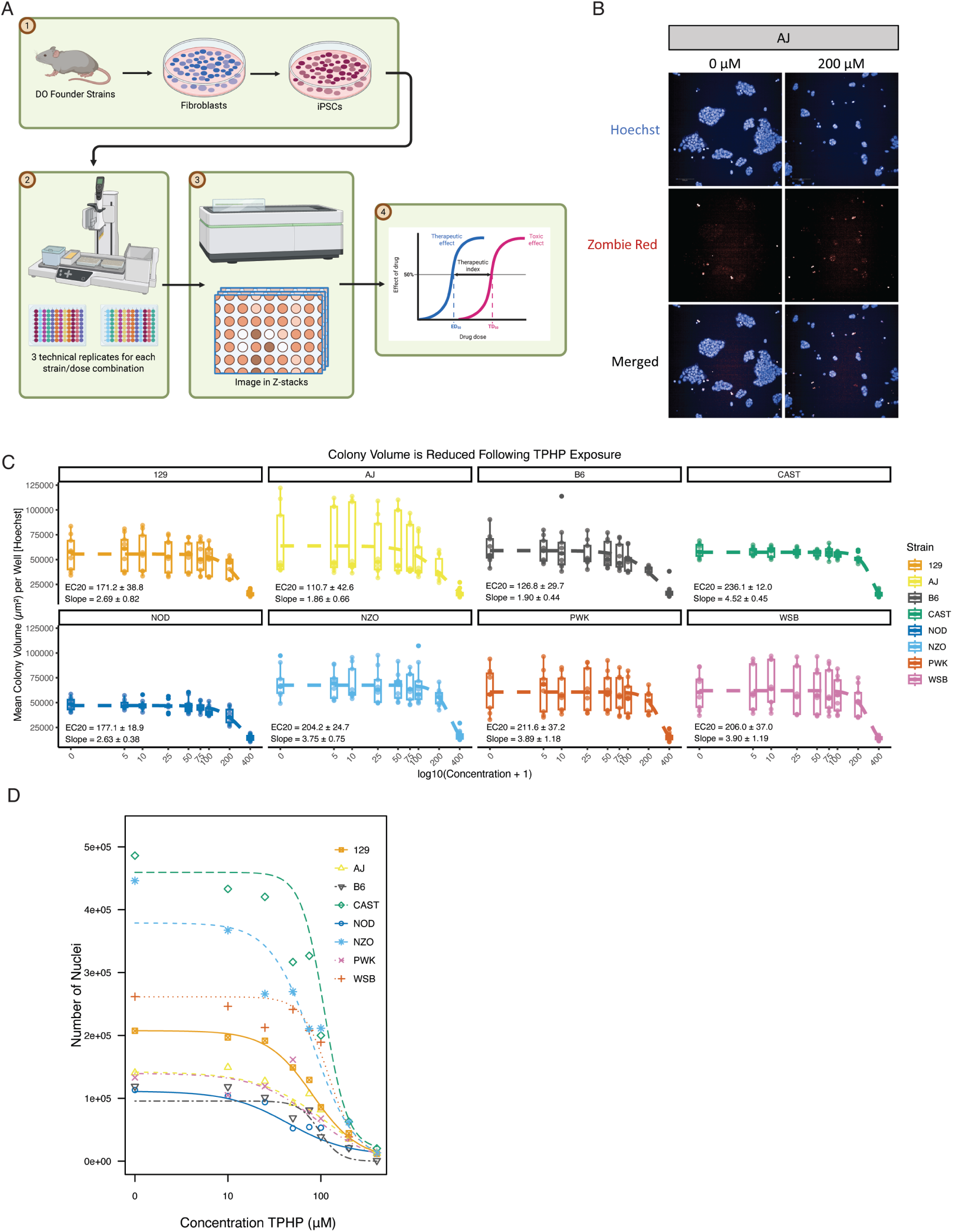
Differences in Inbred DO Founder Exposure Response to TPHP. A) Tail tip fibroblasts were collected from all eight inbred DO founder strains and reprogrammed into iPSCs using a lentiviral vector. iPSCs from each strain were randomly seeded onto 96 well plates (1,500 cells per well, 6 columns per strain). Image analysis was performed on whole well quantifications to generate dose response data for each strain. B) Representative Images from AJ iPSCs showing Hoechst and Zombie Red staining following a 24-hour 0μM or 200μM TPHP exposure C) Mean colony surface area values per well for each founder strain following 24-hour exposure at nine doses (0, 5, 10, 25, 50, 75, 100, 200, and 400μM) of TPHP on 96-well plates (n = 9-18 wells per strain per dose) D) Dose response modeling for each DO founder strain following 24-hour exposure at 8 doses (0, 10, 25, 50, 75, 100, 200, and 400μM) of TPHP. (35,000 cells seeded per well on a 12-well plate)

### DO iPSC lines exhibit continuous variation in TPHP sensitivity

The DO founder experiments were limited to eight inbred genetic backgrounds, but we needed a population with much greater genetic diversity to investigate the effect of genetics on TPHP exposure response. The DO mouse population was derived from the same eight inbred strains, yielding broad phenotypic variation and genetic architecture that supports high resolution genetic mapping. We previously generated a panel of iPSC lines derived from individual DO mice, with each line representing a distinct genetic background suitable for high-throughput *in vitro* genetic analyses (Armstrong et al. 2026). Adequate sample size is required to detect genetic effects. Prior QTL power analyses based on simulated data approximated that a sample size of 200 DO individuals is sufficient to detect loci accounting for greater than 20% of the phenotypic variance with 90% power (Gatti et al. 2014). We used this benchmark to inform sample size and randomly selected 218 DO iPSC lines (106 females, 112 males) for TPHP exposure. For *in vitro* exposure, we sought to identify an effective TPHP concentration corresponding to 20% cytotoxicity (EC20), enabling us to focus on transcriptional and cellular responses in the surviving population after exposure to a fixed dose of TPHP. This concentration is expected to mimic the subset of stem cells that could persist during *in vivo* development following exposure, while also reducing experimental cost and time. To estimate the EC20, we performed dose-response analysis on a subset of ten randomly selected DO iPSC lines using the same eight TPHP doses applied to the founder strains (Figure 2A, 2B). The median dose-response relationship across these lines indicated that a concentration of approximately 80 μM TPHP (0.04% DMSO v/v) produced ∼20% cell loss (Figure 2C). This estimated EC20 acute dose is four to five orders of magnitude higher than most adult human exposure measured to date, and at least 200 times higher than observed embryonic tissue concentrations (Canada 2025; EPA 2016; Zhao et al. 2017), but it is comparable to other *in vitro* TPHP studies that aim at sublethal, mechanistic studies (Feng et al. 2023; Germain and Winn2024; Qi et al. 2019; Wang et al. 2020).

**Figure 2.**
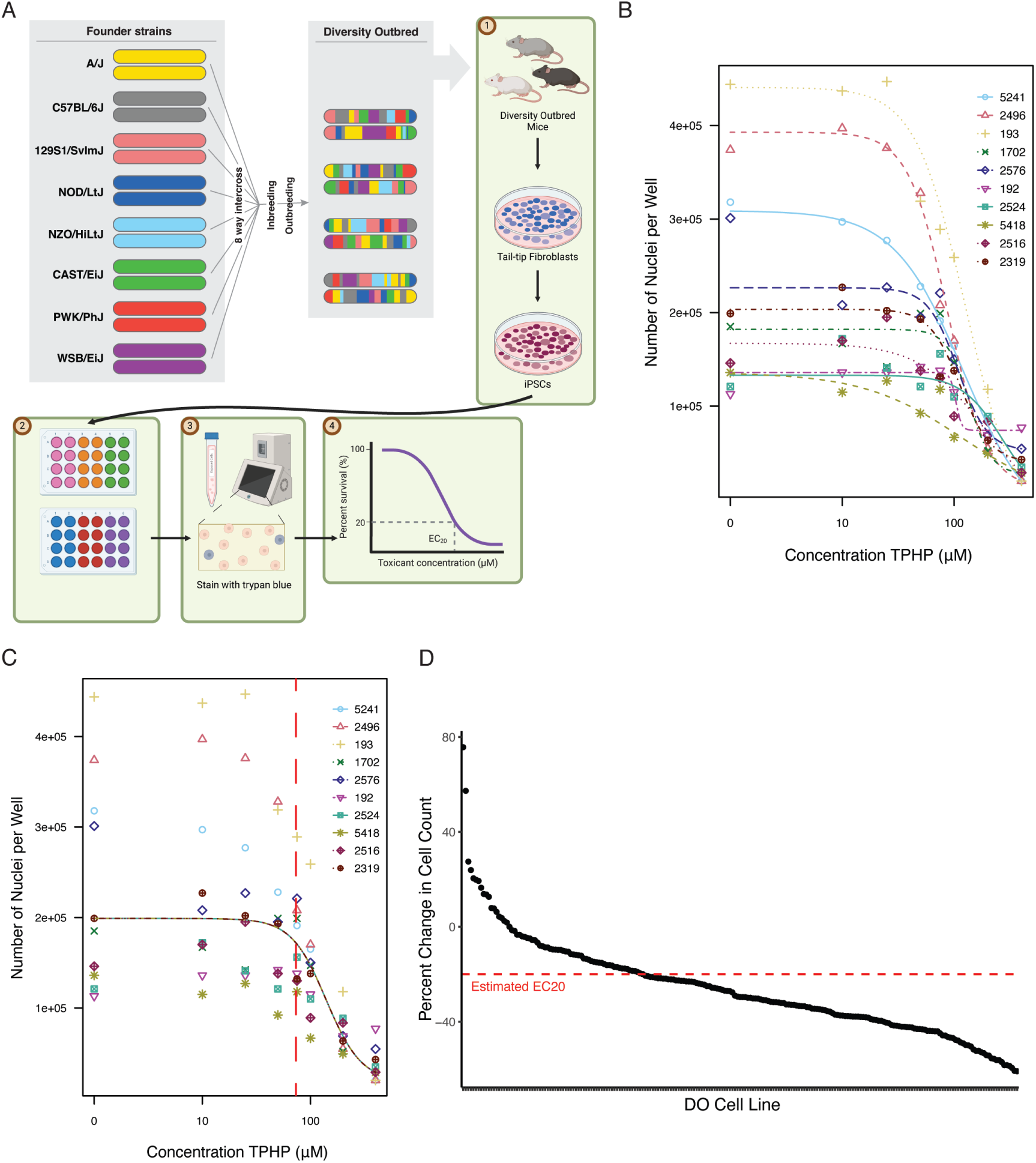
Identification of an Estimated EC20 for TPHP Exposure in DO iPSCs. A) Overview of DO iPSC generation and TPHP EC20 Identification B) Dose response modeling for 10 DO iPSC lines following 24-hour exposure at eight doses (0, 10, 25, 50, 75, 100, 200, and 400μM) of TPHP C) Median dose response modeling for 10 DO iPSC lines, treating all lines as biological replicates to generate a single dose response curve of the population median response. Red dashed line denotes estimated EC20 of 80μM D) Percent Change in Cell Count for 218 DO cell lines following exposure to 80μM TPHP for 24 hours. Red dashed line indicates the estimated EC20 effect of 20% cell loss

We exposed 218 DO iPSC lines to this single EC20 dose of 80 μM TPHP for 24 hours and then collected RNA for RNA sequencing and gene expression analyses. Quantification of cell counts following exposure revealed a continuous distribution of responses across DO iPSC lines (Figure 2D). Most lines exhibited the expected ∼20% reduction in cell number following TPHP exposure; however, we also identified sensitive lines with greater than 40% cell loss as well as highly resistant lines exhibiting potential hormesis (Figure 2D) (Mattson 2008). This continuous and bidirectional variation highlights the extensive phenotypic diversity present within the DO population.

### TPHP disrupts transcriptional programs regulating stem cell proliferation

To identify the molecular mechanisms by which TPHP exposure alters cell number, we performed differential gene expression analysis. A total of 459 genes were significantly upregulated while 845 genes were significantly downregulated in response to TPHP exposure (Figure 3A). To identify overrepresented pathways and gene functions among the differentially expressed genes, we performed gene set enrichment analysis using the Gene Ontology: Biological Process gene set collection published in the Molecular Signatures Database (MSigDB, (Gene Ontology 2026; Liberzon et al. 2011; Subramanian et al. 2005)). Among the significantly suppressed functions were cell cycle and regulation of cell proliferation (Figure 3B). For example, *Ccnd2*, a cyclin that drives G1/S phase cell cycle transition and is essential for stem cell proliferation, was down regulated (37.15% decrease compared to control) following TPHP exposure (Figure 3C) (Zhu et al. 2018). In addition, many FGF and IGF signaling genes also exhibited decreased expression in samples treated with TPHP compared to controls (Figure 3C). Coordinated changes in expression within these growth factor signaling pathways adds to the body of evidence that TPHP disrupts pathways that regulate self-renewal in stem cells (Coutu and Galipeau 2011). These expression changes suggest that decreased cell counts in exposed cells are at least partly due to decreased proliferative capacity. In addition to TPHP-induced shifts in gene expression, we also observed transcriptional heterogeneity across the cell lines. We performed variance component analysis to quantify the sources of this variability, and we found that TPHP exposure (treatment), chromosomal sex of the iPSC line, and genetic background (genotype) differences account for a large portion of expression differences for many genes (Figure 3D).

**Figure 3.**
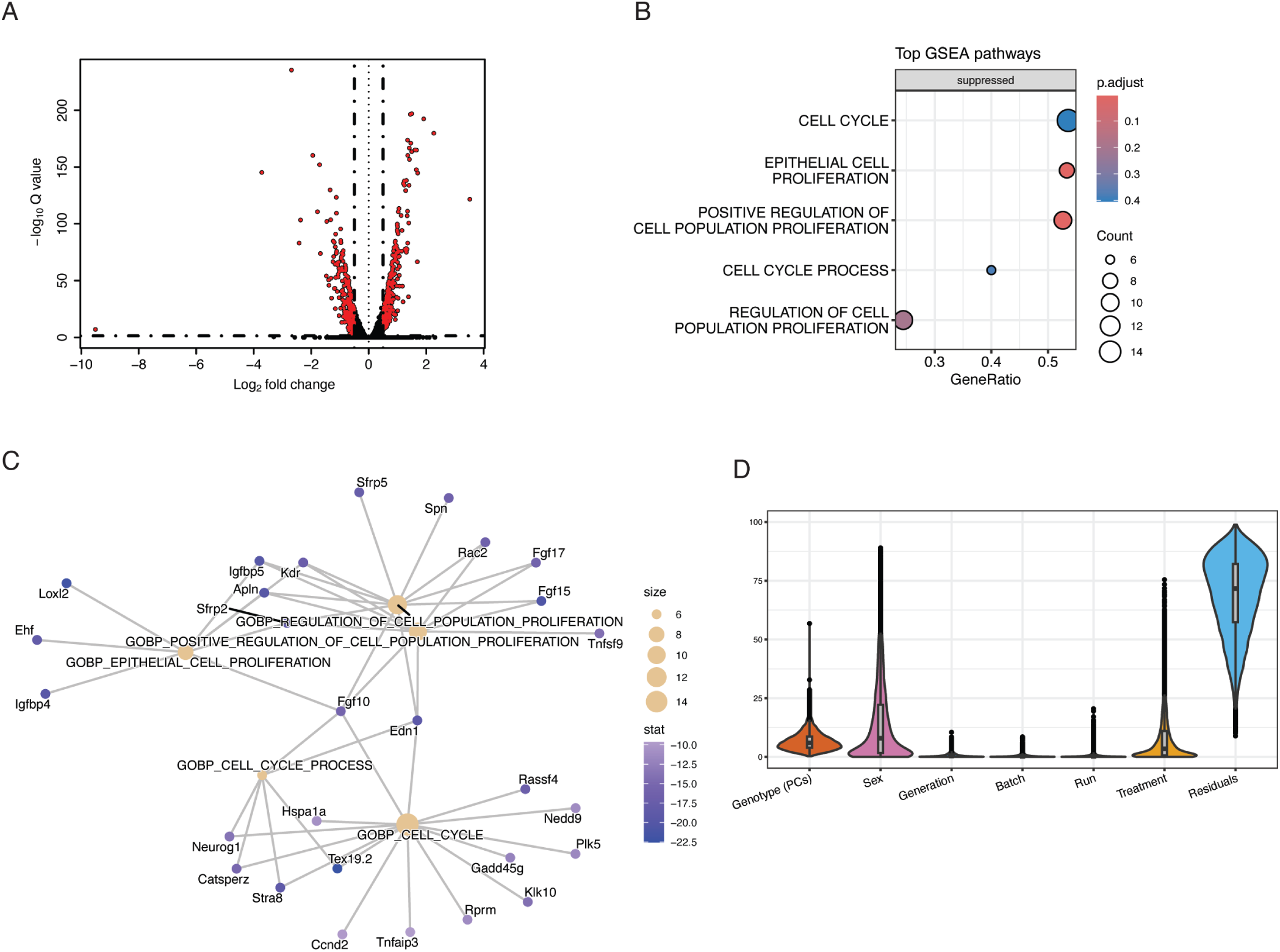
Differential Gene Expression Analysis Following TPHP Exposure. A) Volcano plot of differentially expressed genes between untreated and TPHP exposed genotype-matched samples. Vertical black dashed lines indicate log2-fold change (*LFC*) thresholds at +/-0.5. Red dots are LFC > +/-0.5 with *p*< 0.05 B) Top 5 enriched cell cycle pathways associated with differentially expressed genes between untreated and TPHP exposed genotype-matched samples. C) Gene-concept network of genes within the top 5 enriched cell cycle pathways of differentially expressed genes following TPHP exposure D) Variance component analysis showing the proportion of gene expression variance attributable to genotype, sex, generation, batch, sequencing run, TPHP treatment, and residual effects. Genotype is represented by the first ten principal components derived from genotype probabilities.

### Genetic variation shapes transcriptional responses to TPHP

To further characterize the influence of genetic background on transcriptional responses to TPHP, we performed expression quantitative trait locus (eQTL) mapping. Because genetic variation contributes substantially to differences in gene expression in normal conditions, we needed to distinguish baseline genetic regulatory effects from those specifically induced by TPHP exposure. To remove those baseline genetic effects and focus on loci that affected gene expression specifically in response to TPHP exposure, we mapped eQTLs using the change in gene expression between treated and untreated samples (Caliskan et al. 2015), hereafter referred to as “delta” eQTLs (Figure 4A). This approach allowed us to identify genomic loci associated specifically with exposure-response transcriptional changes. We detected a total of 1463 delta eQTLs (LOD > 8.97) that affect the expression of 1390 genes in response to TPHP exposure. Of these delta eQTLs, 573 are local, meaning the eQTL peak is located close to its target gene and likely acts in *cis*, while 890 delta eQTLs were classified as distal, meaning the eQTL peak is farther away from its target gene and likely acts in *trans* (Figure 4A). Local and distant eQTLs are thought to modulate gene expression through distinct regulatory mechanisms. Local eQTLs typically act in *cis* through direct effects on nearby promoters or enhancers, whereas distant eQTLs often act indirectly in *trans* through intermediate regulators such as transcription factors or signaling molecules (Rockman and Kruglyak 2006). For this reason, we performed gene ontology (GO) term enrichment on local and distant eQTL target genes separately to avoid conflating these mechanistically distinct classes of regulatory variation. TPHP-induced transcripts with local regulatory effects were enriched for pathways related to autophagy, lipid remodeling, and suppressed proliferation and differentiation, while distant eQTL effects were predominantly associated with pathways involved in ribosome biogenesis and steroid hormone signaling (Figure 4B-C).

**Figure 4.**
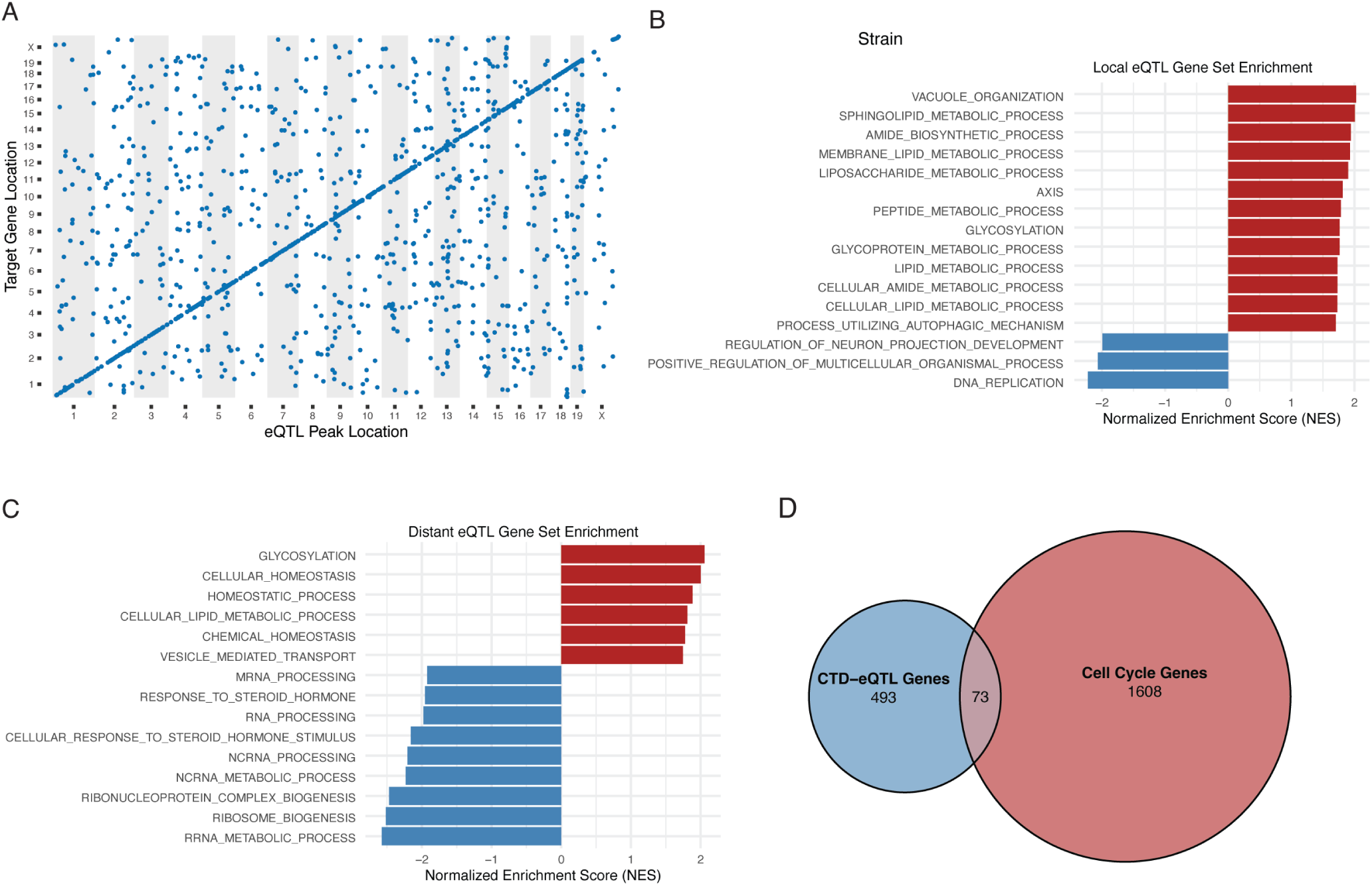
Gene Set Enrichment of TPHP Induced eQTLs Highlight Cell Proliferation Effect. A) Genomic locations of DO iPSC delta eQTLs (LOD > 8.97). Axes display genomic location (numbered by chromosome). Each dot represents a single eQTL. The diagonal line is created by local eQTL dots. All dots off the diagonal line are distant eQTL. B) Gene set enrichment of local eQTL target genes using the gene ontology biological process data set (GO:BP) C) Gene set enrichment of distant eQTL target genes using the gene ontology biological process data set (GO:BP) D) Venn-diagram of shared genes between cell cycle related genes and genes present in both our delta eQTL and CTD associations with TPHP exposure.

### Overlap of TPHP exposure eQTL genes to the Comparative Toxicogenomic Database

To determine whether target genes of TPHP-induced eQTLs have been previously implicated in the toxicological response to TPHP, we intersected our set of 1390 delta eQTL target genes with previously annotated TPHP-interacting genes in the Comparative Toxicogenomic Database (CTD) (Davis et al. 2023). A total of 566 target genes in our delta eQTL dataset were also present in the CTD. Overrepresentation analysis on the overlapping genes between our eQTL target gene set and the TPHP-interacting genes from CTD showed enrichment for metabolism, hormone signaling, and cell surface markers (Supplementary Figure 3). These overlapping genes illustrate a new perspective, that genetic variation regulates these previously annotated TPHP-interacting genes in ways that have not previously been identified.

TPHP has been repeatedly shown to disrupt cell cycle regulation, inducing cell cycle arrest, altering proliferation, and triggering DNA damage responses across multiple model systems, so we further focused our CTD comparison specifically to cell cycle related genes (Qiet al. 2019; Xu et al. 2025). This narrowed focus tests whether our eQTL target genes are enriched for a toxicologically relevant and mechanistically specific subset of TPHP’s transcriptional effects, rather than diluting the signal across the broader, more heterogeneous set of genes CTD associates with TPHP. We tested the 566 gene overlap for enrichment against curated cell-cycle gene sets (MSigDB C5: GO Biological Process, filtered to cell-cycle, proliferation, and stemness related terms) using a one-sided Fisher’s exact test. Of the 566 overlap genes, 73 (12.9%) were annotated to a cell cycle related gene set, compared to 493 overlap genes that are not associated with cell cycle by gene ontology (Figure 4D). This corresponded to a modest enrichment (*OR* = 1.04) but one that did not reach statistical significance (*p* = 0.382). Within this overlap, we observed genes like *Ino80*, which is required for a chromatin remodeler complex that maintains open chromatin architecture at crucial pluripotency gene promoters (Wang et al. 2014). Another overlapping gene, *Csnk2a2*, encodes a catalytic subunit of the CK2 protein kinase complex responsible for upstream regulation of stem cell genes like *Nanog*, *Sox2*, and *Oct4 (*Lu et al. 2014). The 73-cell cycle related genes in the CTD-eQTL overlap highlight that TPHP affects cell cycle across multiple tissues and species.

Despite evidence that TPHP perturbs DNA replication, DNA damage response, and mitotic progression in experimental systems (Qi et al. 2019), these processes were not preferentially enriched among CTD overlapping eQTL genes. We performed a follow-up analysis to test whether the full list of TPHP interacting genes in CTD, pooled across all species in the database, was independently enriched for cell cycle related GO terms. While the CTD-eQTL overlapping gene set was not significantly enriched, we found enrichment for cell cycle related genes (*p* < .0001, *OR* = 3.47) in the full CTD-TPHP gene list (Supplementary Table 2). We next asked whether this signal of cell cycle enrichment within all TPHP associated genes was concentrated within a specific cell cycle mechanism. We defined three independent, literature motivated sub-themes using unmodified GO:BP term definitions from MSigDB: DNA repair/replication, spindle/chromosome segregation, significant enrichment for all three themes, comparison against a null distribution of size-matched, curated GO:BP gene sets revealed that much of this enrichment is consistent with general annotation and study bias shared between curated pathway databases and the CTD chemical-gene interaction database. Enrichment for DNA repair/replication and spindle/chromosome segregation fell within the range typical of unrelated curated pathways (*OR* 95th percentile threshold = 8.4 across 40 control terms), while G1/S checkpoint (*OR* = 11.0) exceeded the observed range of controls, suggesting a comparatively stronger signal for that theme (Supplementary Table 3).

### Delta eQTL hotspots

Within eQTL data we can identify regulatory hotspots, genomic locations that are associated with the variation in expression of many target genes simultaneously. To identify regulatory hotspots in our delta eQTL data, we relaxed our inclusion criteria to include suggestive eQTLs (LOD > 7.42) and then quantified the density of distant eQTLs across the genome, yielding 23 hotspots with at least 5 target genes each (Figure 5A, Supplementary Table 4). The most prominent hotspots were found on Chromosomes 13 and 15 and regulated the expression of 118 and 65 target genes, respectively. Targets of the Chr 13 hotspot include genes associated with transcription and protein phosphorylation, while targets of the Chr 15 hotspot include genes with roles in cell cycle and differentiation. To identify which genes within these eQTL hotspots likely explain, or “mediate”, the observed variation in expression of target genes, we applied mediation analysis (Chick et al. 2016). The Chr 13 hotspot contains numerous closely related KRAB zinc-finger transcription factors as well as computationally annotated gene models, and mediation analysis was unable to identify a single best mediator for the Chr 13 hotspot (Supplementary Figure 4). Mediation of the Chr 15 hotspot identified the LIF receptor gene *Lifr* as the strongest candidate mediator (Figure 5B). *Lifr* is a known regulator of self-renewal that was previously identified in eQTL studies of DO ESCs and in untreated DO iPSCs (Armstrong et al. 2026; Skelly et al. 2020). We then performed haplotype effect analysis to characterize the effect of genotype within the chromosome 15 hotspot interval on the expression of the hotspot’s 65 target genes. This revealed a 3:5 founder allele pattern, with haplotype effects from the three more recently wild-derived strains (CAST/EiJ, PWK/PhJ, and WSB/EiJ) clustering together (Figure 5C-D). This same 3:5 haplotype pattern was observed for the *Lifr* hotspot effect in untreated iPSCs, further supporting the conclusion that this interval is a major genetic regulator of cell proliferation and cell-cycle that drives additional gene expression variability in response to TPHP exposure (Graf et al. 2011). The repertoire of target genes in the TPHP induced hotspot differs somewhat from those found in untreated iPSCs. Notably, following exposure to TPHP, cell lines carrying wild-derived alleles in the hotspot region showed higher expression of genes related to lipid and membrane metabolism and vesicle trafficking (*Rai14, Washc1, Dtna, Mtmr12, Acsbg2, Akr1cl*) compared to cell lines with alleles from classically inbred strains. Inversely, lines with alleles from classically inbred strains showed higher expression of genome-stability and developmental genes (*Fancd2, Runx1t1, Zscan4-ps3, Foxp1, Qser1, Lefty1, Sfrp2, Fgf17*). Two additional pluripotency-associated genes (*Firre, Khdc3*) were more highly expressed in lines with the wild-derived alleles, suggesting that the two haplotype groups may employ distinct mechanisms for maintaining pluripotency and genome stability in the context of TPHP exposure.

**Figure 5.**
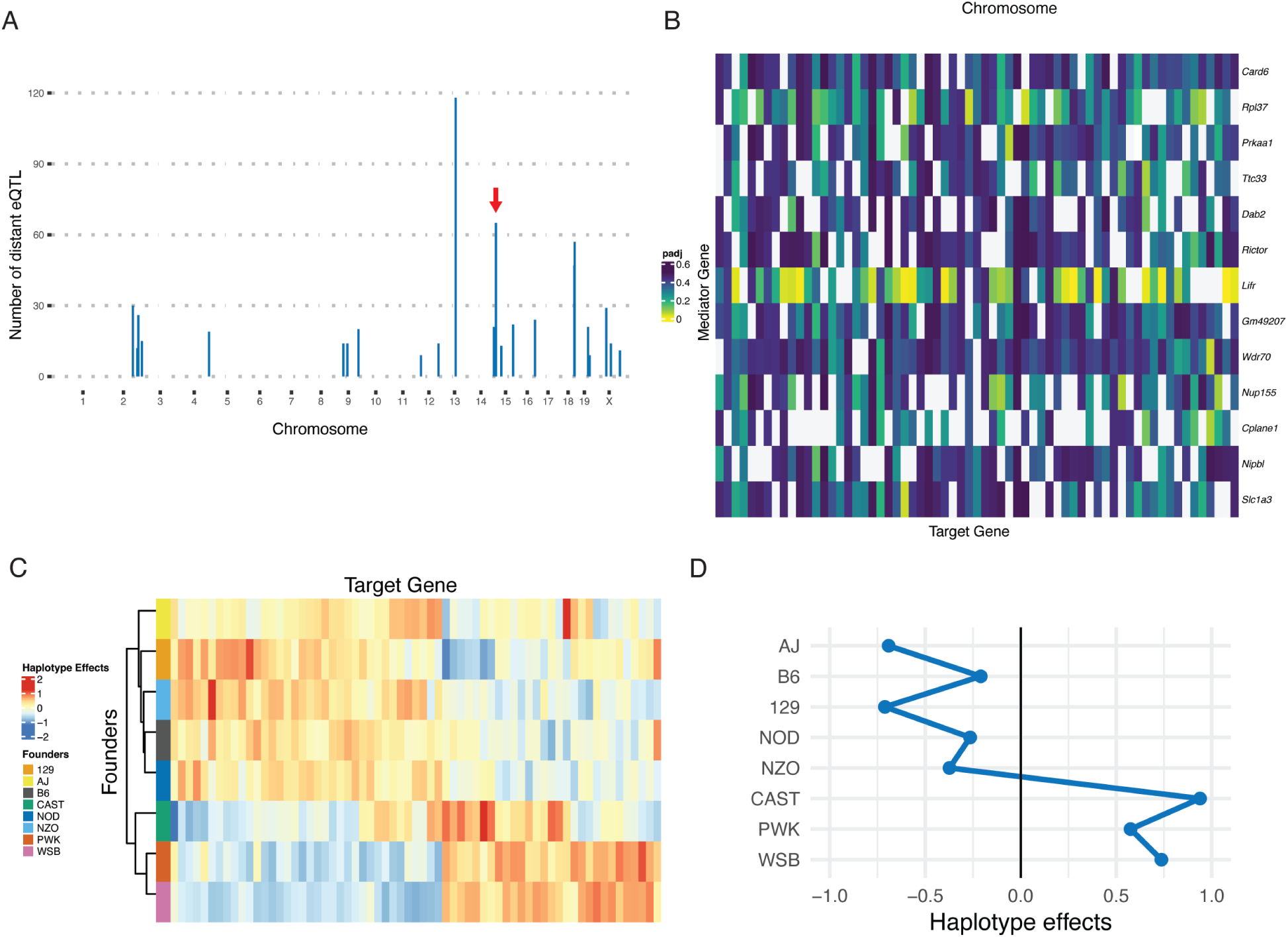
TPHP Induced eQTL Hotspot on Chromosome 15 Affects *Lifr* Regulatory Pathway. A) Histogram showing colocalization of distant delta eQTLs to genomic hotspots. Suggestive and greater distant eQTLs (LOD > 7.42) were summed across a sliding window of 50 chromosome-based markers with 10 marker shift and totals were plotted (y-axis) according to their genome location (x-axis). The Chr 15 hotspot is denoted by a red arrow. B) Heat map of mediation analysis identifying *Lifr* as the strongest mediator for the chromosome 15 hotspot. Potential mediators were identified using the change in LOD score and the associated adjusted p-values for each mediator-target interaction are represented by color. C) Heatmap of the haplotype effects at the 65 distant eQTL within the chromosome 15 hotspot showing a 3:5 split between the wild-derived and classic inbred strain alleles. D) Haplotype effect at the local eQTL for *Lifr* showing a 3:5 split, with more recently wild-derived strains grouping separately from the classic inbred strains.

### TPHP induces strain-dependent changes in cell cycle progression

To better understand how TPHP induced gene expression changes alter important stem cell functions, we investigated the effect of TPHP on stem cell cycle. Unlike somatic cells, pluripotent stem cells utilize cyclins, CDKs, and related core cell cycle components to maintain a truncated G1 phase, which is thought to limit the window during which differentiation signals can act on the cell. This contrasts with adult tissue-specific stem cells, which instead tend to sit in a quiescent G0 state that restricts differentiation (Liu et al. 2019). We utilized flow cytometry to quantify G1/S/G2 cell cycle ratios in the eight inbred DO founder strain iPSCs following TPHP exposure. Cells exposed to TPHP had reduced populations of cells in G2 and increased populations of cells in G1, although the magnitude of these changes differed between strains (Figure 6A-B). These differences were statistically significant with some genetic backgrounds showing larger shifts in cell cycle phase ratios than others upon exposure to TPHP and G1/S checkpoint regulation.

**Figure 6.**
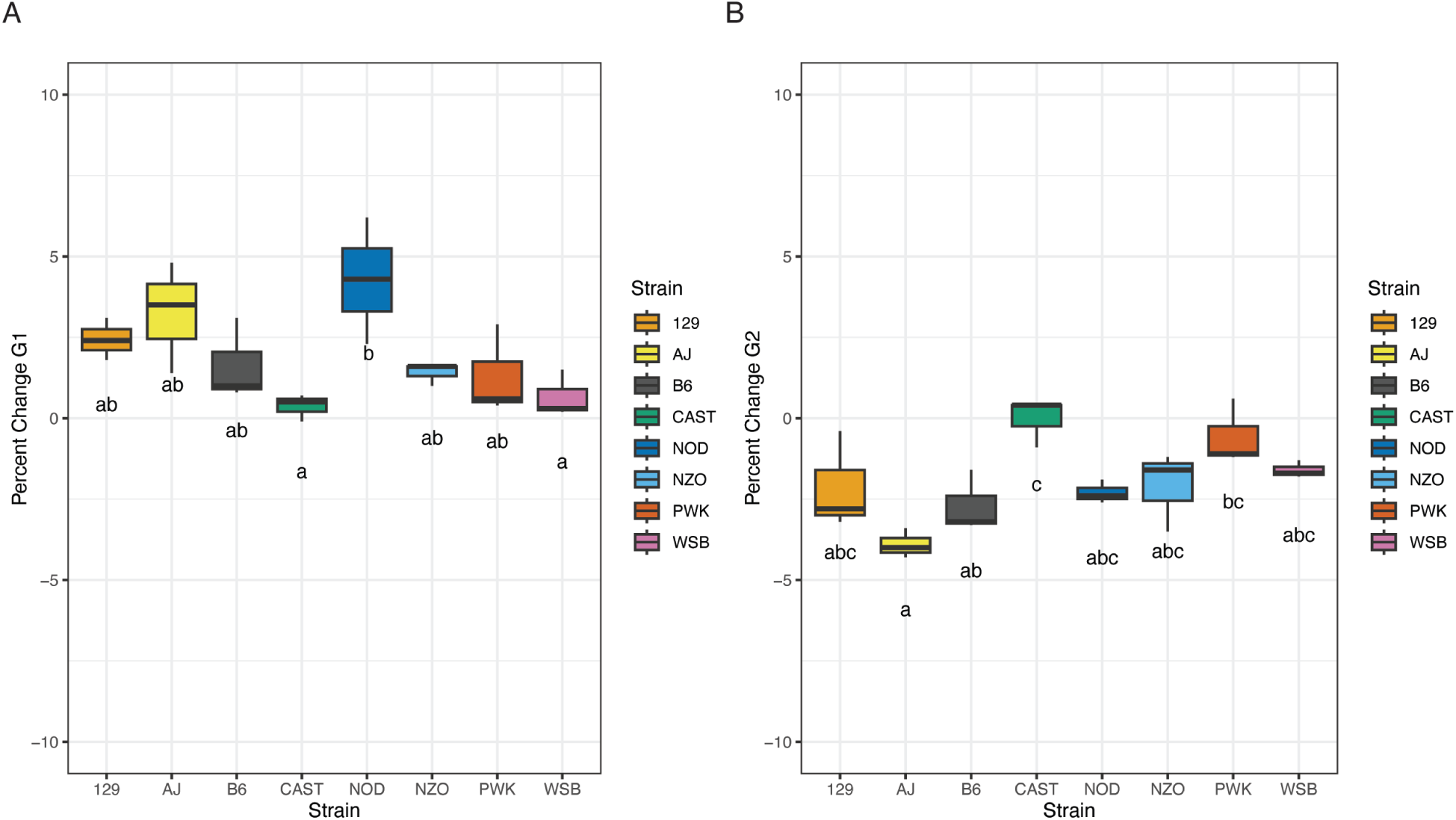
TPHP Exposure Inhibits Stem Cell Cycle Progression. A) Percent change in the number of cells in G1 of cell cycle following 24-hour exposure to 60μM TPHP (DO founder EC20). Letters indicate ANOVA bins, where shared letters indicate similarity while no shared letters indicate significant difference. B) Percent change in the number of cells in G2 of cell cycle following 24-hour exposure to 60μM TPHP (inbred DO founder median EC20). Letters indicate ANOVA bins, where shared letters indicate similarity while no shared letters indicate significant difference.

## DISCUSSION

Our findings establish that genetic background is a primary determinant of cellular response to TPHP exposure in a genetically diverse population of iPSCs. Across the eight DO inbred founder strains, we observed heritable differences in colony morphology, cell death, and proliferation following TPHP exposure. This variation extended continuously across the larger panel of 218 DO iPSC lines, with hypersensitive lines losing more than 40% of cells and other lines showing no appreciable loss, and sometimes a modest gain, in cell cumber consistent with hormesis. A wide range of response was observed between the DO iPSC lines and potentially indicates that a substantial portion of what is often treated as “response variability” in TPHP studies could be genetically programmed and polygenic (Evans and Johnson 2001; Haber et al.2002; Lewis et al. 2017). This has a direct implication for how chemical safety is currently assessed. Models built on a single genetic background, as is standard practice in most *in vitro* toxicology, are structurally unable to detect this level of variation and may therefore systematically underestimate the range of human sensitivity to TPHP exposure (Abdo et al.2015; French et al. 2015).

The transcriptional changes observed in exposed cells point to a specific mechanism where TPHP disrupts the signaling networks that maintain stem cell self-renewal and proliferation. Downregulation of *Ccnd2*, a cyclin required for G1/S transition in pluripotent cells, together with coordinated suppression of FGF and IGF signaling genes, suggests that TPHP interferes directly with the growth factor signaling stem cells use to maintain a proliferative, undifferentiated state. This is consistent with the phenotypic shift we observed toward G1 and away from G2 across founder strains. These changes raise a developmentally relevant possibility, where if TPHP exposure blunts self-renewal signaling in pluripotent cells *in vivo*, similar disruption during early embryogenesis could shift the timing or efficacy of early developmental transitions.

Our eQTL analysis adds a genetic foundation to this mechanistic picture. The contrasting pathways linked to local versus distant eQTL effects point to two somewhat different modes by which TPHP exposure reshapes the transcriptome. Transcripts under local regulatory control were enriched for autophagy and lipid remodeling pathways (Figure 5C, top), a pattern that fits with a direct cellular response to chemical stress common when a xenobiotic like TPHP disrupts membrane integrity or metabolic balance. Local effects were also enriched for suppression of pathways tied to proliferation and differentiation, including DNA replication and neuron projection development, which indicate a stress response that comes at the expense of normal growth programs. Distant eQTL effects highlight a different but concurrent response, with enrichment concentrated in ribosome biogenesis and steroid hormone signaling (Figure 5C, bottom). Because these genes are not physically linked to their regulatory variants, this likely reflects a more indirect route of influence, where downstream consequences of TPHP exposure ripple outward through broader regulatory networks rather than acting locally. Taken together, these results suggest that TPHP disrupts stem cell homeostasis on two fronts, a proximal stress response tied to autophagy and lipid handling, and through more diffuse trans-regulatory effects on translation and endocrine signaling.

When we intersected our response eQTL target genes with those independently annotated as TPHP interacting in the Comparative Toxicogenomic Database, we found 566 genes in common between the two datasets. Our study adds 824 new genes that have not previously been associated with TPHP response and provides additional evidence that genetic variation modulates the response of these genes in a diverse cell population. The overlap between the eQTL and CTD datasets (566 genes) showed only a modest, non-significant enrichment for cell cycle and stemness gene sets (*OR* = 1.04, *p* = 0.382). However, when we examined the full set of TPHP-associated genes curated in CTD across all species, cell cycle terms were strongly enriched (*OR* = 3.47, *p* < .0001), and this signal was not uniform across cell cycle sub-mechanisms: G1/S checkpoint regulation specifically exceeded the range expected from general curation overlap between pathway databases (*OR* = 11.0), whereas enrichment for DNA repair/replication and spindle/chromosome segregation fell within the range attributable to shared annotation bias, derived from CTD annotations being accumulated from the literature.

The distal regulatory hotspot on chromosome 15 is driven by *Lifr*, a gene previously identified as a distal regulator of proliferation in both DO embryonic stem cells and untreated DO iPSCs (Armstrong et al. 2026; Skelly et al. 2020). The fact that this same locus re-emerges as a driver of TPHP induced transcriptional change strengthens the case that *Lifr* sits at a uniquely regulatory node in stem cell proliferative control, one that is both constitutively active and alterable by chemical exposure. In humans, alterations in *LIFR* expression are associated with cancer progression, infertility and implantation failure, and a variety of abnormalities in skeletal and muscle development, exhibiting how LIFR sits at a signaling axis that is used across multiple tissues (Braun et al. 2025; Halder et al. 2022; Zutautas et al. 2023). The haplotype effects we observed at this locus are also notable; the three wild-derived founders (CAST/EiJ, PWK/PhJ, WSB/EiJ) responded differently than the five classical inbred strains, with wild-derived lines upregulating lipid, membrane and vesicle trafficking genes following exposure, while classical strains upregulated genome stability and developmental patterning genes. Because several of these genes, *Lefty1* and *Sfrp2* in particular, function in early embryonic patterning and left-right axis specification, this divergence suggests genetic background influences both the level and the composition of TPHP response genes. The additional upregulation of the pluripotency-associated genes *Firre* and *Khdc3* specifically in wild-derived strains further suggests these two haplotype groups may utilize distinct routes to preserving pluripotency and genomic stability under chemical stress.

Together, this suggests that while TPHP affects various aspects cell cycle, like DNA repair/replication and spindle/chromosome segregation, the effects are disproportionately concentrated at the G1/S transition, consistent with our direct observation of a shift in cell cycle towards G1 in some strain backgrounds and with *Ccnd2* downregulation at the transcriptional level. The absence of a comparably strong cell-cycle signal in the eQTL-CTD overlap may reflect the gene regulatory architecture of TPHP response in the DO population. Local delta-eQTL were enriched for pathways associated with reduced proliferation, indicating that some gene regulatory effects act directly through genes involved in these processes. But the bulk of the effects may act through broader regulatory pathways, as illustrated by the *Lifr* trans eQTL hotspot and enrichment of steroid hormone signaling among distal eQTL genes.

Overall, these results extend our understanding of TPHP toxicity in stem cells as a genotype-by-environment phenomenon, supporting future consideration of genetic background in toxicological response for chemicals like TPHP. This has practical implications for risk assessment, where EC20 or similar threshold doses – while useful for standardizing exposure screens – may substantially misrepresent risk for individuals or subpopulations at the extreme ends of the exposure response distribution. Our study is limited to an *in vitro* pluripotent stem cell population model, and the extent to which this gene regulatory architecture is conserved in differentiated tissue types or during later developmental windows remains untested. Future work incorporating differentiation into specific lineages, longitudinal exposure studies, or functional validation of candidate eQTL genes, including *Lifr* and the G1/S checkpoint genes, will help clarify whether the regulatory hotspots identified here represent generalizable determinants of chemical susceptibility or are specific to the pluripotent state. Allele matched DO or Collaborative Cross mice for *Lifr* or other genomic loci of interest provide a potential validation platform to verify our *in vitro* findings are maintained *in vivo*. Nonetheless, this work demonstrates that genetically diverse cell panels such as the DO iPSC resource offer a tractable and scalable approach for identifying the genetic determinants of chemical toxicity, a capability that single genotype models cannot provide.

## ACKNOWLEDGEMENTS

We gratefully acknowledge the contribution of Genome Technologies and Protein Sciences Services at The Jackson Laboratory for expert assistance and services in support of the work described in this publication. We sincerely appreciate the assistance of Lacie Low for protocol refinement and preliminary data collection. Vectors were a gift from Dr. Matthias Stadtfeld.

## DATA AVAILABILITY

GigaMUGA genotyping data, raw and processed files, and code are available on Figshare (10.6084/m9.figshare.34007223). RNA-seq data are available from GEO GSE 333520.

## AUTHOR CONTRIBUTIONS

**Madison Armstrong**, methodology, formal analysis, investigation, writing-original draft, visualization, software; **Kathryn Janeczko**, investigation, data curation; **Hannah B. Dewey**, methodology, software; **Anne Czechanski**, methodology, data curation; **Qiongyu Chen**, methodology, data curation; **Emily Swanzey**, methodology; **Callan O’Connor**, methodology; **Whitney Martin**, methodology, resources; **Selcan Aydin**, methodology, software; **Steven C. Munger**, conceptualization, funding acquisition, methodology, formal analysis, supervision, writing – review & editing; **Laura Reinholdt**, conceptualization, funding acquisition, formal analysis, supervision, writing – review & editing

