## Supplementary Figure 1 for "The Flame Retardant Triphenyl Phosphate Induces Varying Responses in Genetically Diverse Mouse Induced Pluripotent Stem Cells"

A

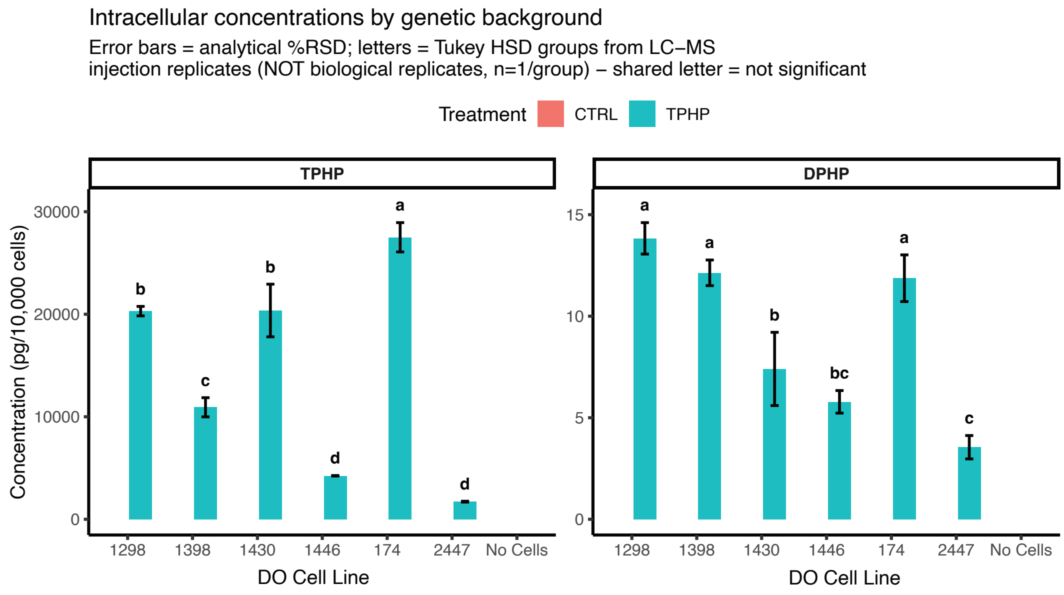


B

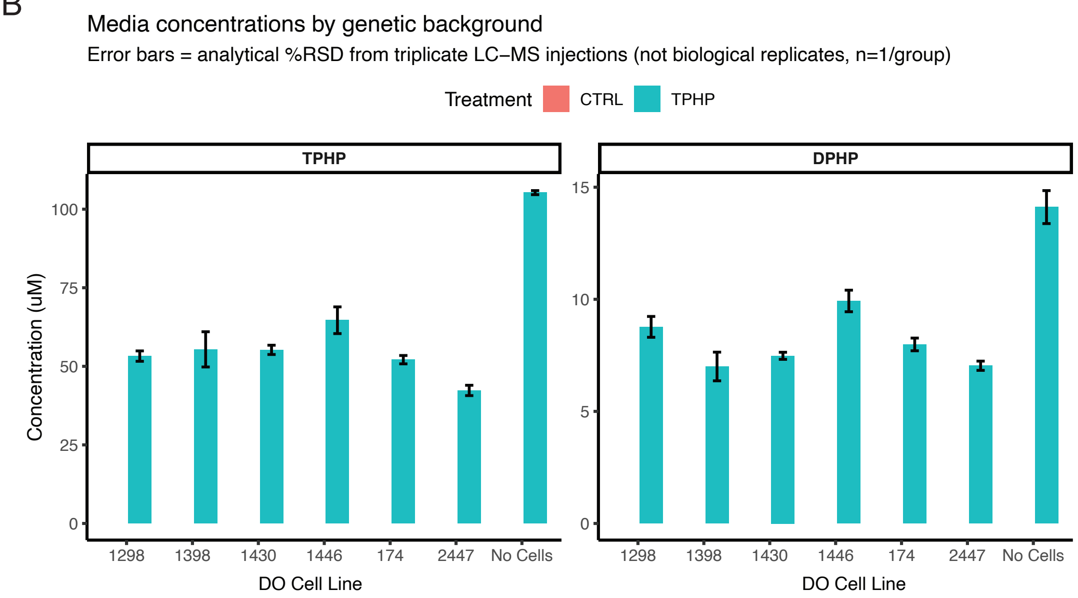


**Supplementary Figure 1.** A) Bar plots of TPHP and its metabolite DPHP intracellular concentrations (pg/10,000 cells). ANOVA binning is represented by letters, distinct letters indicate significance. Control samples had no detectable TPHP or DPHP. A no cell sample (media only) was also included as a negative control. B) Bar plots of TPHP and its metabolite DPHP media concentrations (µM). Control samples had no detectable TPHP or DPHP. A no cell sample (media only) was also included as a TPHP positive control and to test for spontenous TPHP degreadation into DPHP.
