## Supplementary Figure 2 for "The Flame Retardant Triphenyl Phosphate Induces Varying Responses in Genetically Diverse Mouse Induced Pluripotent Stem Cells"

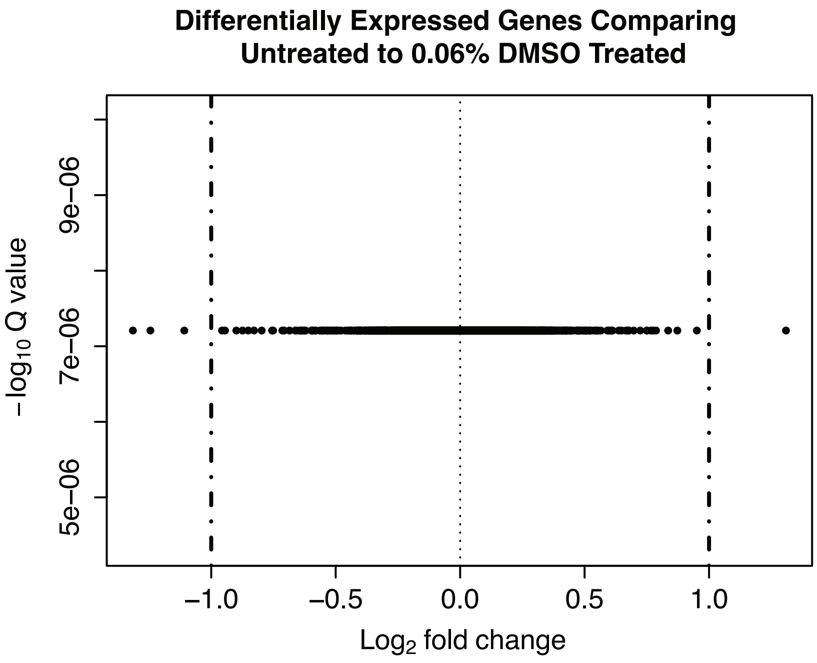


**Supplementary Figure 2.** Volcano plot of differentially expressed genes between untreated and 0.06% DMSO exposed genotype-matched samples. Vertical black dashed lines indicate log2-fold change (LFC) threshold sat +/- 1.0.
