## Supplementary Figure 3 for "The Flame Retardant Triphenyl Phosphate Induces Varying Responses in Genetically Diverse Mouse Induced Pluripotent Stem Cells"

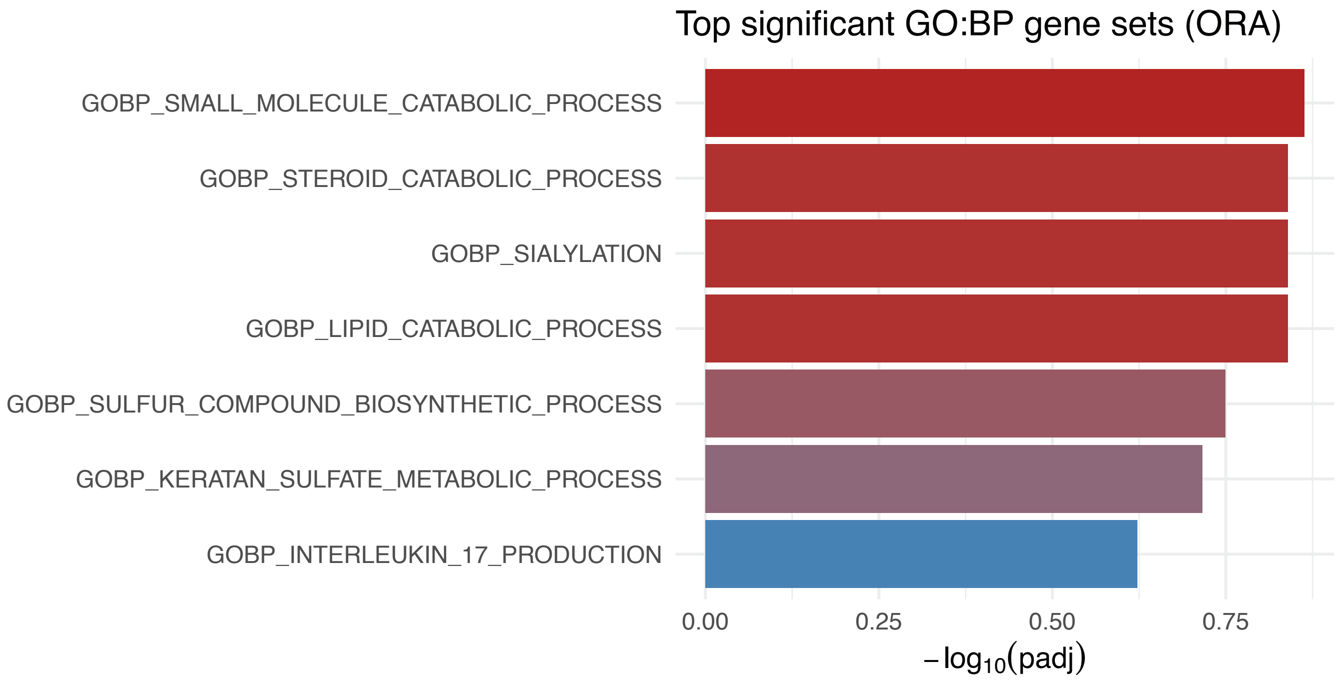


**Supplementary Figure 3.** Overrepresentation analysis on the overlapping genes between our delta eQTL target gene set and the CTD-TPHP gene list (padj < 0.25).
