## Supplementary Figure 4 for "The Flame Retardant Triphenyl Phosphate Induces Varying Responses in Genetically Diverse Mouse Induced Pluripotent Stem Cells"

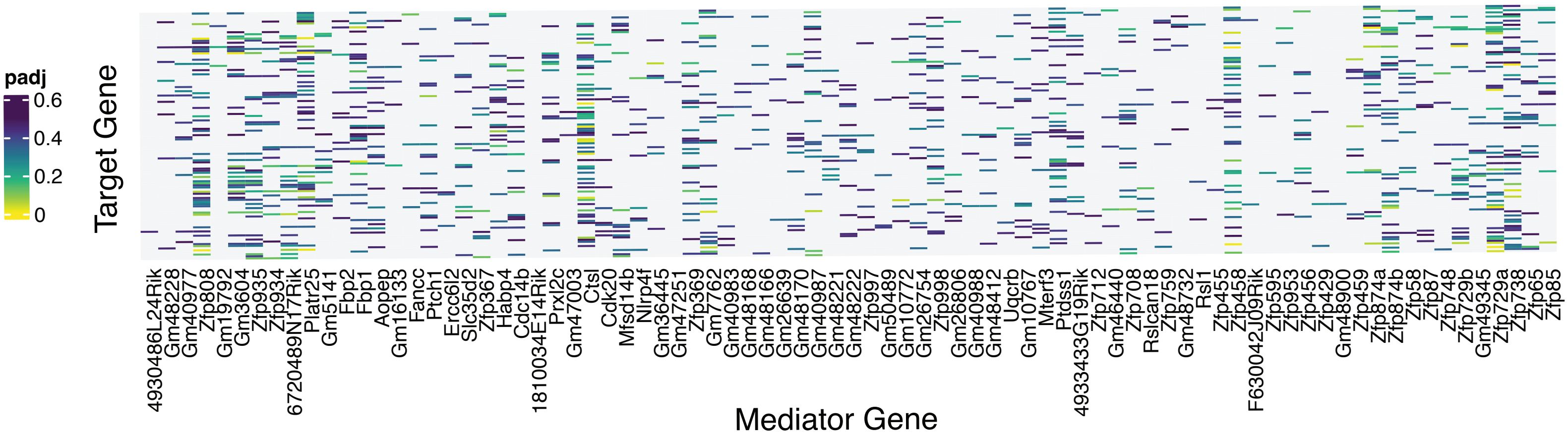


**Supplementary Figure 4.** Heat map of mediation analysis in the Chr 13 hotspot. Potential mediators were identified using the change in LOD score and the associated adjusted p-values for each mediator-target interaction are represented by color.
