## Supplementary Table 1 for "The Flame Retardant Triphenyl Phosphate Induces Varying Responses in Genetically Diverse Mouse Induced Pluripotent Stem Cells"

| variable | mean | sd | mean_percent | sd_percent |
|---|---|---|---|---|
| Genotype (PCs) | 0.07 | 0.04 | 6.74 | 3.91 |
| Sex | 0.15 | 0.17 | 14.55 | 16.78 |
| Generation | 0.01 | 0.01 | 0.72 | 0.95 |
| Batch | 0.01 | 0.01 | 0.5 | 0.71 |
| Run | 0.01 | 0.01 | 0.72 | 1.31 |
| Treatment | 0.08 | 0.11 | 8.22 | 11.24 |
| Residuals | 0.69 | 0.17 | 68.54 | 17.05 |

**Supplementary Table 1.** Data table with variance component analysis results showing the proportion of gene expression variance attributable to genotype, sex, generation, batch, sequencing run, TPHP treatment, and residual effects. Genotype is represented by the first ten principal components derived from genotype probabilities.
