## Supplementary Table 2 for "The Flame Retardant Triphenyl Phosphate Induces Varying Responses in Genetically Diverse Mouse Induced Pluripotent Stem Cells"

| Category | Cell cycle | Not cell cycle |
| --- | --- | --- |
| In CTD-TPHP list | 1688 | 6661 |
| Not in CTD-TPHP | 1574 | 18648 |

| Statistic | Value |
| --- | --- |
| p-value | 2.8273477372529e-18 |
| Odds ratio | 3.0020258370617 |
| Alternative hypothes | greater |

**Supplementary Table 2.** (Upper) Contingency table of genes in CTD-TPHP that are associated with GO cell cycle terms compared to a universe of all CTD annotated genes across all species. (Lower) Fisher's exact test (one-sided) results for cell cycle enrichment in CTD-TPHP gene set.
