## Supplementary Table 3 for "The Flame Retardant Triphenyl Phosphate Induces Varying Responses in Genetically Diverse Mouse Induced Pluripotent Stem Cells"

| theme | p | OR | in_ctd_in_theme | in_ctd_not_in_theme | in_theme_not_in_ctd | not_in_theme_not_in_ctd | padj |
|---|---|---|---|---|---|---|---|
| dna_repair_replication | 5.0352E-150 | 7.107125 | 461 | 10013 | 306 | 47239 | 1.5106E-149 |
| spindle_segregation | 2.7644E-122 | 8.317287 | 340 | 10134 | 191 | 47354 | 4.1465E-122 |
| g1s_checkpoint | 9.86116E-81 | 10.99943 | 193 | 10281 | 81 | 47464 | 9.86116E-81 |

**Supplementary Table 3.** Contingency table of genes in CTD-TPHP that are associated with GO cell cycle sub-mechanism terms compared to a universe of all CTD annotated genes across all species. Fisher's exact test (one-sided) results for cell cycle sub-mechansim enrichment in CTD-TPHP gene set.
