## Supplementary Table 4 for "The Flame Retardant Triphenyl Phosphate Induces Varying Responses in Genetically Diverse Mouse Induced Pluripotent Stem Cells"

| seqnames | start | end | width | strand | distant_rna | distant_rna_sugg | chr |
|---|---|---|---|---|---|---|---|
| 2 | 133137580 | 135369384 | 2231805 | * | 6 | 30 | 2 |
| 2 | 154222167 | 155175199 | 953033 | * | 4 | 12 | 2 |
| 2 | 159266282 | 160022749 | 756468 | * | 4 | 26 | 2 |
| 2 | 174308507 | 178750740 | 4442234 | * | 5 | 15 | 2 |
| 4 | 144517808 | 148658966 | 4141159 | * | 8 | 19 | 4 |
| 9 | 37657992 | 40290208 | 2632217 | * | 4 | 14 | 9 |
| 9 | 57245307 | 58161751 | 916445 | * | 4 | 14 | 9 |
| 9 | 107854248 | 110046068 | 2191821 | * | 5 | 20 | 9 |
| 12 | 21381711 | 25277424 | 3895714 | * | 4 | 9 | 12 |
| 12 | 104305432 | 105274067 | 968636 | * | 4 | 14 | 12 |
| 13 | 60922092 | 67916067 | 6993976 | * | 21 | 118 | 13 |
| 14 | 121105663 | 123282432 | 2176770 | * | 7 | 21 | 14 |
| 15 | 4903055 | 9087988 | 4184934 | * | 20 | 65 | 15 |
| 15 | 31506242 | 32332659 | 826418 | * | 4 | 13 | 15 |
| 15 | 86167813 | 87094559 | 926747 | * | 5 | 22 | 15 |
| 16 | 84638503 | 85603942 | 965440 | * | 4 | 24 | 16 |
| 18 | 73560283 | 75175950 | 1615668 | * | 5 | 47 | 18 |
| 18 | 75225201 | 76523127 | 1297927 | * | 8 | 57 | 18 |
| 19 | 46891604 | 48040610 | 1149007 | * | 4 | 21 | 19 |
| 19 | 54145583 | 55214529 | 1068947 | * | 4 | 9 | 19 |
| X | 69835597 | 72454926 | 2619330 | * | 5 | 29 | X |
| X | 91546064 | 94136692 | 2590629 | * | 5 | 14 | X |
| X | 133961254 | 135568990 | 1607737 | * | 4 | 11 | X |

**Supplementary Table 4.** Data table of distant eQTL hotspots identified by QTLretrievR, where distant is defined as eQTL greater than 2Mbp from its target gene. Distant_rna is distant eQTL LOD > 8.97, distant_rna_sugg is distant eQTL suggLOD > 7.42. The suggLOD values were used for hotspot analysis. Start, end, and width are genomic locations in base pair (bp).
